# Type I interferon susceptibility distinguishes SARS-CoV-2 from SARS-CoV

**DOI:** 10.1101/2020.03.07.982264

**Authors:** Kumari G. Lokugamage, Adam Hage, Maren de Vries, Ana M. Valero-Jimenez, Craig Schindewolf, Meike Dittmann, Ricardo Rajsbaum, Vineet D. Menachery

## Abstract

SARS-CoV-2, a novel coronavirus (CoV) that causes COVID-19, has recently emerged causing an ongoing outbreak of viral pneumonia around the world. While distinct from SARS-CoV, both group 2B CoVs share similar genome organization, origins to bat CoVs, and an arsenal of immune antagonists. In this report, we evaluate type-I interferon (IFN-I) sensitivity of SARS-CoV-2 relative to the original SARS-CoV. Our results indicate that while SARS-CoV-2 maintains similar viral replication to SARS-CoV, the novel CoV is much more sensitive to IFN-I. In Vero and in Calu3 cells, SARS-CoV-2 is substantially attenuated in the context of IFN-I pretreatment, while SARS-CoV is not. In line with these findings, SARS-CoV-2 fails to counteract phosphorylation of STAT1 and expression of ISG proteins, while SARS-CoV is able to suppress both. Comparing SARS-CoV-2 and influenza A virus in human airway epithelial cultures (HAEC), we observe the absence of IFN-I stimulation by SARS-CoV-2 alone, but detect failure to counteract STAT1 phosphorylation upon IFN-I pretreatment resulting in near ablation of SARS-CoV-2 infection. Next, we evaluated IFN-I treatment post infection and found SARS-CoV-2 was sensitive even after establishing infection. Finally, we examined homology between SARS-CoV and SARS-CoV-2 in viral proteins shown to be interferon antagonists. The absence of an equivalent open reading frame (ORF) 3b and changes to ORF6 suggest the two key IFN-I antagonists may not maintain equivalent function in SARS-CoV-2. Together, the results identify key differences in susceptibility to IFN-I responses between SARS-CoV and SARS-CoV-2 that may help inform disease progression, treatment options, and animal model development.

**Importance:** With the ongoing outbreak of COVID-19, differences between SARS-CoV-2 and the original SARS-CoV could be leveraged to inform disease progression and eventual treatment options. In addition, these findings could have key implications for animal model development as well as further research into how SARS-CoV-2 modulates the type I IFN response early during infection.

**Article Summary:** SARS-CoV-2 has similar replication kinetics to SARS-CoV, but demonstrates significant sensitivity to type I interferon treatment.

## Introduction

At the end of 2019, a cluster of patients in Hubei Province, China was diagnosed with a viral pneumonia of unknown origins. With community links to the Huanan seafood market in Wuhan, the disease cluster had echoes of the severe acute respiratory syndrome coronavirus (SARS-CoV) outbreak that emerged at the beginning of the century (1). The 2019 etiologic agent was identified as a novel coronavirus, 2019-nCoV, and subsequently renamed SARS-CoV-2 (2). The new virus has nearly 80% nucleotide identity to the original SARS-CoV and the corresponding CoV disease, COVID-19, has many of the hallmarks of SARS-CoV disease including fever, breathing difficulty, bilateral lung infiltration, and death in the most extreme cases (3, 4). In addition, the most severe SARS-CoV-2 disease corresponded to old age (>50 years old), health status, and healthcare workers, similar to both SARS- and MERS-CoV (5). Together, the results indicate SARS-CoV-2 infection and disease have strong similarity to the original SARS-CoV epidemic occurring nearly two decades earlier.

In the wake of the outbreak, major research efforts have sought to rapidly characterize the novel CoV to aid in treatment and control. Initial modeling studies predicted (6) and subsequent cell culture studies confirmed that spike protein of SARS-CoV-2 utilizes human angiotensin converting enzyme 2 (ACE2) for entry, the same receptor as SARS-CoV (7, 8). Extensive case studies indicated a similar range of disease onset and severe symptoms seen with SARS-CoV (5). Notably, less severe SARS-CoV-2 cases have also been observed and were not captured in the original SARS-CoV outbreak. Importantly, screening and treatment guidance has relied on previous CoV data generated with SARS-CoV and MERS-CoV. Treatments with both protease inhibitors and type-I interferon (IFN-I) have been employed (4); similarly, remdesivir, a drug targeting viral polymerases, has been reported to have efficacy against SARS-CoV-2 similar to findings with both SARS- and MERS-CoV (9-12). Importantly, several vaccine efforts have been initiated with a focus on the SARS-CoV-2 spike protein as the major antigenic determinant (13). Together, the similarities with SARS-CoV have been useful in responding to the newest CoV outbreak.

The host innate immune response is initiated when viral products are recognized by host cell pattern recognition receptors, including Toll-like receptors (TLRs) and RIG-I-like receptors (RLRs) (14, 15). This response ultimately results in production of IFN-I and other cytokines, which together are essential for an effective antiviral response (16). IFN-I then triggers its own signaling cascade via its receptor, in an autocrine or paracrine manner, which induces phosphorylation of signal transducers and activators of transcription 1 (STAT1) and STAT2. Together, STAT1, STAT2, and a third transcription factor, IRF9, form the Interferon Stimulated Gene Factor 3 (ISGF3) complex, which is essential for induction of many IFN-stimulated genes (ISGs), and ultimately elicit an effective antiviral response (17, 18). To establish productive replication, viruses have developed different mechanisms to escape this antiviral response targeting different parts of the IFN-I response machinery (19).

In this study, we further characterize SARS-CoV-2 and compare it to the original SARS-CoV. Using Vero E6 cells, we demonstrate that SARS-CoV-2 maintains similar viral replication kinetics as SARS-CoV following a low dose infection. In contrast, we find that SARS-CoV-2 is significantly more sensitive to IFN-I pretreatment as compared to SARS-CoV. Infection of IFN-I competent Calu3 2B4 cells resulted in reduced SARS-CoV-2 replication compared to SARS-CoV. Similar to Vero cells, Calu3 cells pretreated with IFN-I had a greater reduction of replication of SARS-CoV-2 compared to SARS-CoV. In human airway epithelial cultures, SARS-CoV-2 showed robust replication and an absence of IFN-I stimulation contrasting influenza A virus. However, pretreatment with IFN-I confirmed SARS-CoV-2 sensitivity and inability to control IFN-I responses once initiated. These results suggest distinct changes between SARS-CoV and SARS-CoV-2 in terms of IFN-I antagonism and we subsequently examined sequence homology between the SARS-CoV and SARS-CoV-2 viral proteins that may be responsible for these differences. Together, the results suggest SARS-CoV-2 lacks the same capacity to control the IFN-I response as SARS-CoV.

## Results

### SARS-CoV-2 is sensitive to IFN-I pre-treatment

Our initial studies infected Vero E6 cells using a low multiplicity of infection (MOI) to explore the viral replication kinetics of SARS-CoV-2 relative to SARS-CoV. Following infection, we found that both SARS-CoV and SARS-CoV-2 replicate with similar kinetics, peaking 48 hours post infection (**Fig. 1A**). While SARS-CoV-2 titer was slightly lower than that of SARS-CoV at 24 hours post infection, the results were not statistically different. By 48 hours, replication of both viruses had plateaued and significant cytopathic effect (CPE) was observed for both SARS-CoV and SARS-CoV-2 infections. Together, the results indicated that SARS-CoV and SARS-CoV-2 replicate with similar replication kinetics in Vero E6 cells.

**Figure 1.**
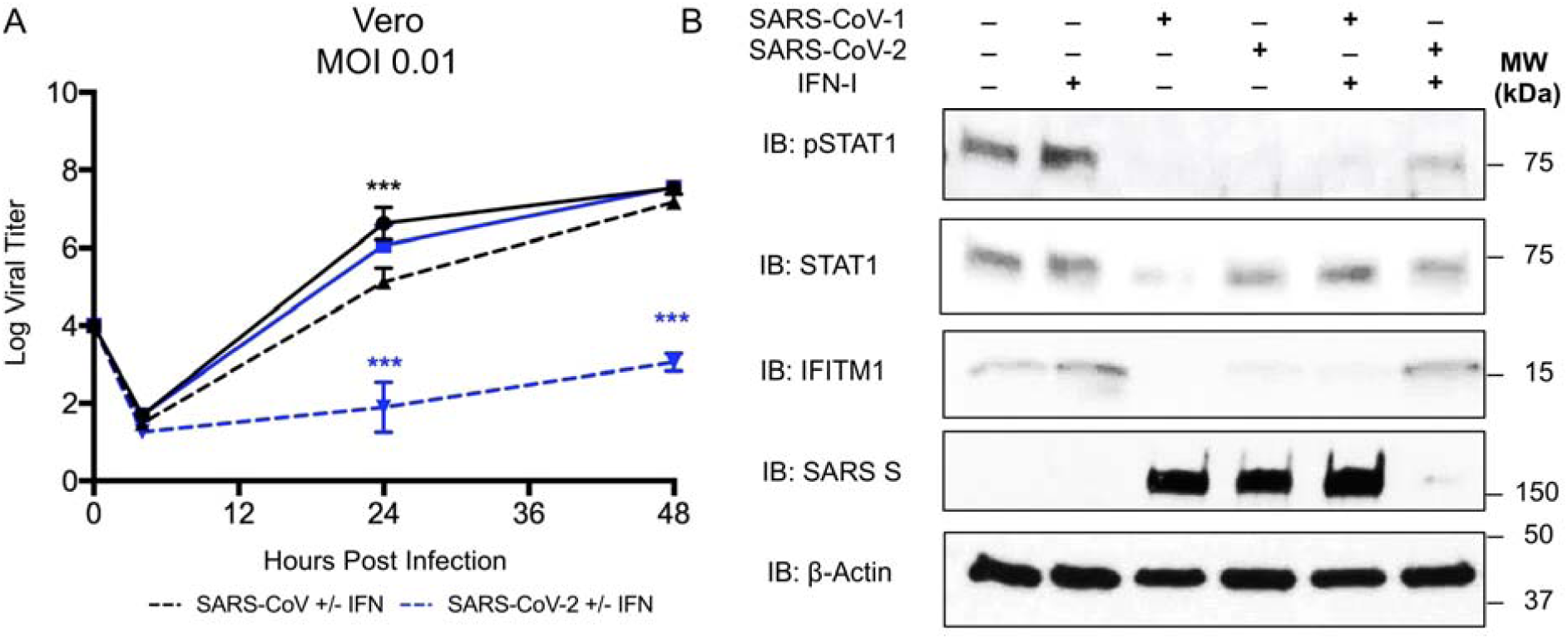
SARS-CoV-2 sensitive to type I IFN pretreatment. A) Vero E6 cells were treated with 1000 U/mL recombinant type I (hashed line) IFN or mock (solid line) for 18 hours prior to infection. Cells were subsequently infected with either SARS-CoV WT (black) or SARS-CoV-2 (blue) at an MOI of 0.01 as described above. Each point on the line graph represents the group mean, N ≥ 3. All error bars represent SD. The two tailed student’s t-test was used to determine P-values: *** P < 0.001. B) Vero cell protein lysates from IFN-I treated and untreated cells were probed 48 hours post infection by Western blotting for phosphorylated STAT1 (Y701), STAT1, IFITM1, SARS spike, and Actin.

We next evaluated the susceptibility of SARS-CoV-2 to IFN-I pretreatment. Treatment with IFN-I (recombinant IFN-α) has been attempted as an antiviral approach for a wide variety of pathogens including hepatitis B and C viruses as well as HIV (20). During both the SARS and MERS-CoV outbreaks, IFN-I has been employed with limited effect (21, 22). In this study, we pretreated Vero E6 cells with 1000 U/mL of recombinant IFN-I (IFN-α) 18 hours prior to infection. Vero E6 lack the capacity to produce IFN-I, but are able to respond to exogenous treatment (23). Following pretreatment with IFN-I, SARS-CoV infection has a modest reduction in viral titer of 1.5 log_10_ plaque forming units (PFU) as compared to untreated control 24 hours post infection (**Fig. 1A**). However, by 48 hours, SARS-CoV has nearly equivalent viral yields as the untreated conditions (7.2 log_10_PFU versus 7.5 log_10_PFU). In contrast, SARS-CoV-2 shows a significant reduction in viral replication following IFN-I treatment. At both 24 and 48 hours post infection, SARS-CoV-2 had massive 3-log_10_ (24 HPI) and 4-log_10_ (48 HPI) drops in viral titer as compared to control untreated cells. Together, the results demonstrate a clear sensitivity to a primed IFN-I response in SARS-CoV-2, which is not observed with SARS-CoV.

To explore differences in IFN-I antagonism between SARS-CoV and SARS-CoV-2, we examined both STAT1 activation and IFN stimulated gene (ISG) expression following IFN-I pretreatment and infection. Examining Vero cell protein lysates, we found that IFN-I treated cells infected with SARS-CoV-2 induced phosphorylated STAT1 by 48 hours post infection (**Fig. 1B**). SARS-CoV had no evidence of STAT1 phosphorylation in either IFN-I treated or untreated cells, illustrating robust control over IFN-I signaling pathways. In contrast, SARS-CoV-2 is unable to control signaling upon IFN-I treatment. Examining further, IFITM1, a known ISG (17), had increased protein expression in the context of SARS-CoV-2 infection following IFN-I pretreatment compared to SARS-CoV under the same conditions (**Fig. 1B**). Basal STAT1 levels are reduced during SARS-CoV infection relative to uninfected control and to a lesser extent during SARS-CoV-2, likely due to the mRNA targeting activity of non-structural protein 1 (NSP1) (24). However, IFN-I treatment results in augmented protein levels for IFITM1 following SARS-CoV-2 infection as compared to untreated SARS-CoV-2. In contrast, IFN-I treated SARS-CoV had no significant increase in IFITM1 relative to control infection. Together, the STAT1 phosphorylation, ISG production, and viral protein levels indicate that SARS-CoV-2 lacks the same capacity to modulate the IFN-I stimulated response as the original SARS-CoV.

### SARS-CoV-2 attenuated in interferon competent cells

While capable of responding to exogenous IFN-I, Vero cells lack the capacity to produce IFN-I following infection which likely plays a role in supporting robust replication of a wide range of viruses. To evaluate SARS-CoV-2 in a IFN-I responsive cell type, we infected Calu3 2B4 cells, a lung epithelial cell line sorted for ACE2 expression and previously used in coronavirus and influenza research (25). Using an MOI of 1, we examined the viral replication kinetics of SARS-CoV-2 relative to SARS-CoV in Calu3 cells. We found that both SARS-CoV and SARS-CoV-2 replicate with similar overall kinetics, peaking 24 hours post infection (**Fig. 2A**). However, SARS-CoV-2 replication is slightly attenuated relative to SARS-CoV at 24 hours post infection (0.82 log_10_ reduction). The attenuation in viral replication expands at 48 hours (1.4 log_10_ reduction) indicating a significant change in total viral titers between SARS-CoV and SARS-CoV-2. Notably, no similar attenuation was observed in untreated Vero cells (**Fig. 1A**) suggesting possible immune modulation of SARS-CoV-2 infection in the respiratory cell line, due to differential sensitivity to secreted IFN-I during infection. We next evaluated the susceptibility of SARS-CoV-2 to IFN-I pretreatment in Calu3 cells. When pretreating cells with 1000 U/mL of recombinant IFN-I 18 hours prior to infection, SARS-CoV infection has a modest reduction in viral titer of ∼0.8 log_10_ PFU as compared to untreated control at both 24 and 48 hours post infection (**Fig. 2A**). Similar to Vero cell results, SARS-CoV-2 shows a significant reduction in viral replication following IFN-I treatment in Calu3 cells. At both 24 and 48 hours post infection, SARS-CoV-2 had 2.65-log_10_ (24 HPI) and 2-log_10_ (48 HPI) drops in viral titer, respectively, as compared to control untreated Calu3 cells. Together, the results demonstrate a clear sensitivity to a primed IFN-I response in SARS-CoV-2, which is not observed with SARS-CoV.

**Figure 2.**
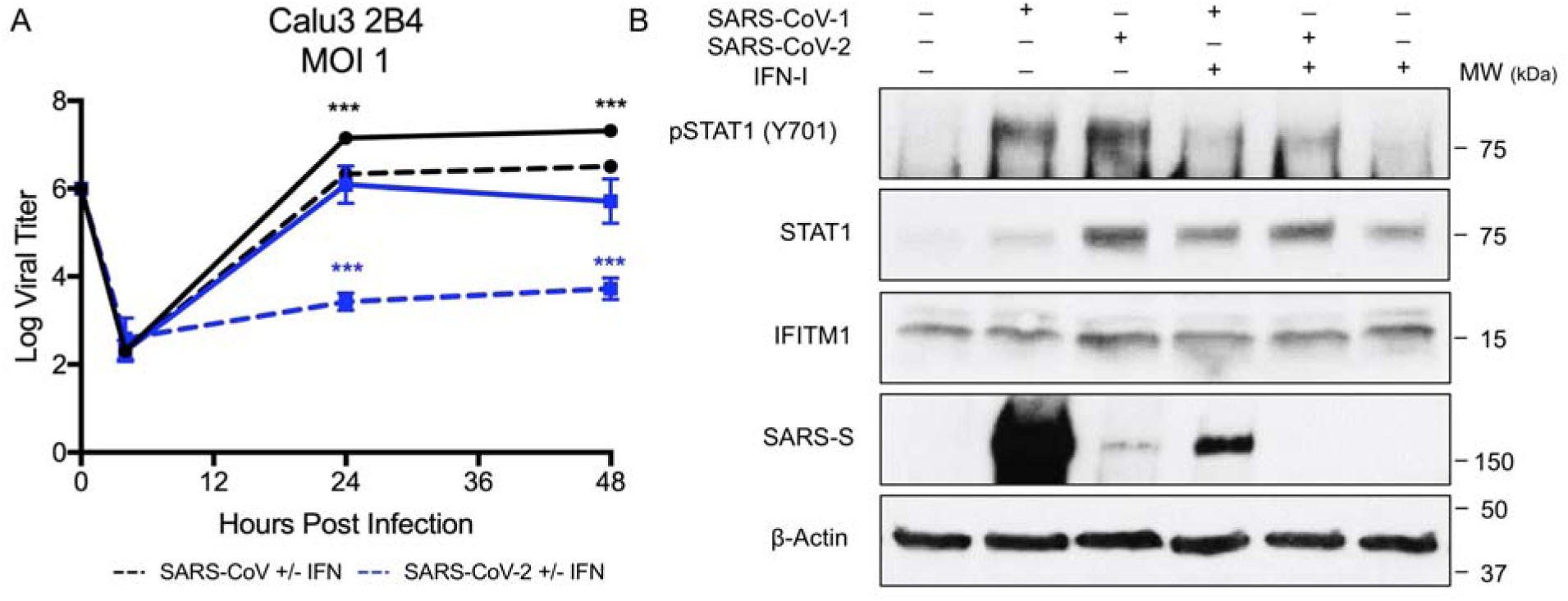
SARS-CoV-2 attenuated and IFN-I sensitive in Calu3 respiratory cells. A) Calu3 2B4 cells were treated with 1000 U/mL recombinant type I (hashed line) IFN or mock (solid line) for 18 hours prior to infection. Cells were subsequently infected with either SARS-CoV WT (black) or SARS-CoV-2 (blue) at an MOI of 1. Each point on the line graph represents the group mean, N ≥ 3. All error bars represent SD. The two tailed student’s t-test was used to determine P-values: *** P < 0.001. B) Calu3 cell protein lysates from IFN-I treated and untreated cells were probed 48 hours post infection by Western blotting for phosphorylated STAT1 (Y701), STAT1, IFITM1, SARS spike, and Actin.

To further evaluate activation by IFN-I, we examined both STAT1 phosphorylation and ISG expression following infection of Calu3 2B4 cells at 48 hours. Probing cell protein lysates, we found that untreated cells infected with SARS-CoV or SARS-CoV-2 induced phosphorylated STAT1 by 48 hours post infection (**Fig. 2B**). However, the level of STAT1 is markedly diminished in SARS-CoV infection as compared to SARS-CoV-2 suggesting the original epidemic strain disrupts the expression of the ISG. The diminished STAT1 levels in SARS-CoV correspond to robust spike expression in untreated cells. In contrast, SARS-CoV-2 spike is reduced as compared to SARS-CoV, consistent with lower replication observed in untreated Calu3 cells (**Fig. 2A**). The noticeable IFITM1 expression levels observed in untreated Calu3 cells infected with SARS-CoV-2 that were not observed in Vero cells may signify higher ISG levels in Calu3 cells during SARS-CoV-2 infection and may contribute to its attenuation relative to SARS-CoV.

Following IFN-I pretreatment in Calu3 cells the differences in STAT1 activation and ISG induction are less prominent compared the Vero experiments, likely due to an already induced IFN-I response to the virus infection. However, we still found a reduction in STAT1 phosphorylation and ISG induction in IFN-I pretreated SARS-CoV versus SARS-CoV-2 infection. Total STAT1 levels were augmented in IFN-I treated cells following SARS-CoV infection to values similar to control IFN-I treated cells. In contrast, SARS-CoV-2 STAT1 levels remained amplified relative to SARS-CoV, but were similar to untreated SARS-CoV-2 infection. Although marginal, IFITM1 expression was slightly increased in IFN-I pretreatment SARS-CoV-2 relative to SARS-CoV. Importantly, the spike protein levels show a significant impact of IFN-I on SARS-CoV infection consistent with titer. For SARS-CoV-2, the spike blot demonstrates the massive attenuation in the presence of IFN-I. Overall, the results in Calu3 cells are consistent with Vero cell findings and indicate a significant sensitivity of SARS-CoV-2 to IFN-I pretreatment.

### SARS-CoV-2 blocks IFN-I signaling in polarized human airway epithelial cultures

Having established IFN-I sensitivity in Vero cells and Calu3 respiratory cell lines, we next sought to evaluate SARS-CoV-2 in polarized human airway epithelial cultures (HAEC). These cultures provide both the complexity of different cell types and the architecture of the human respiratory epithelium. Previous work with SARS-CoV had already established its capacity to control the IFN-I response in human airway cultures (26). Therefore, to examine the impact on overall level of IFN-I signaling in HAECs, we compared SARS-CoV-2 infection to influenza A virus (H1N1) infection, known to induce robust innate immune responses in these cultures. Briefly, HAEC were pretreated with IFN-α on the basolateral side prior to and during infection (**Fig. 3A**). Apical washes were collected at multiple time points and analyzed for viral infectious titers by plaque assay. In addition, culture lysates at endpoint were examined for STAT1 phosphorylation by western blot. Both influenza A and SARS-CoV-2 triggered different levels of immune stimulation (**Fig. 3B**). Influenza A virus infection alone induced robust expression of both total and phosphorylated STAT1 48 hours post infection. In stark contrast, SARS-CoV-2 infection alone indicated no increase in STAT1 or phosphorylated STAT1. IFN-I pretreatment resulted in robust induction of STAT1 and pSTAT1 in all cultures. Despite the distinct immune stimulation profiles upon infection alone, both viruses robustly replicated in the HAEC, with influenza A virus achieving higher viral yields as compared to SARS-CoV-2 (**Fig. 3C & 3D**). Pretreatment reduced influenza infection ∼1.5 log_10_ at both 24 and 48 hours post infection, consistent with previous reports (27). In contrast, IFN-I pretreatment nearly ablated SARS-CoV-2 with 2-log_10_ and 4-log_10_ reduction in viral titers at 24 and 48 hours relative to untreated controls. Together, the results highlight the capacity of SARS-CoV-2 to prevent IFN-I signaling, but also demonstrate the sensitivity of SARS-CoV-2 to IFN-I pretreatment.

**Figure 3.**
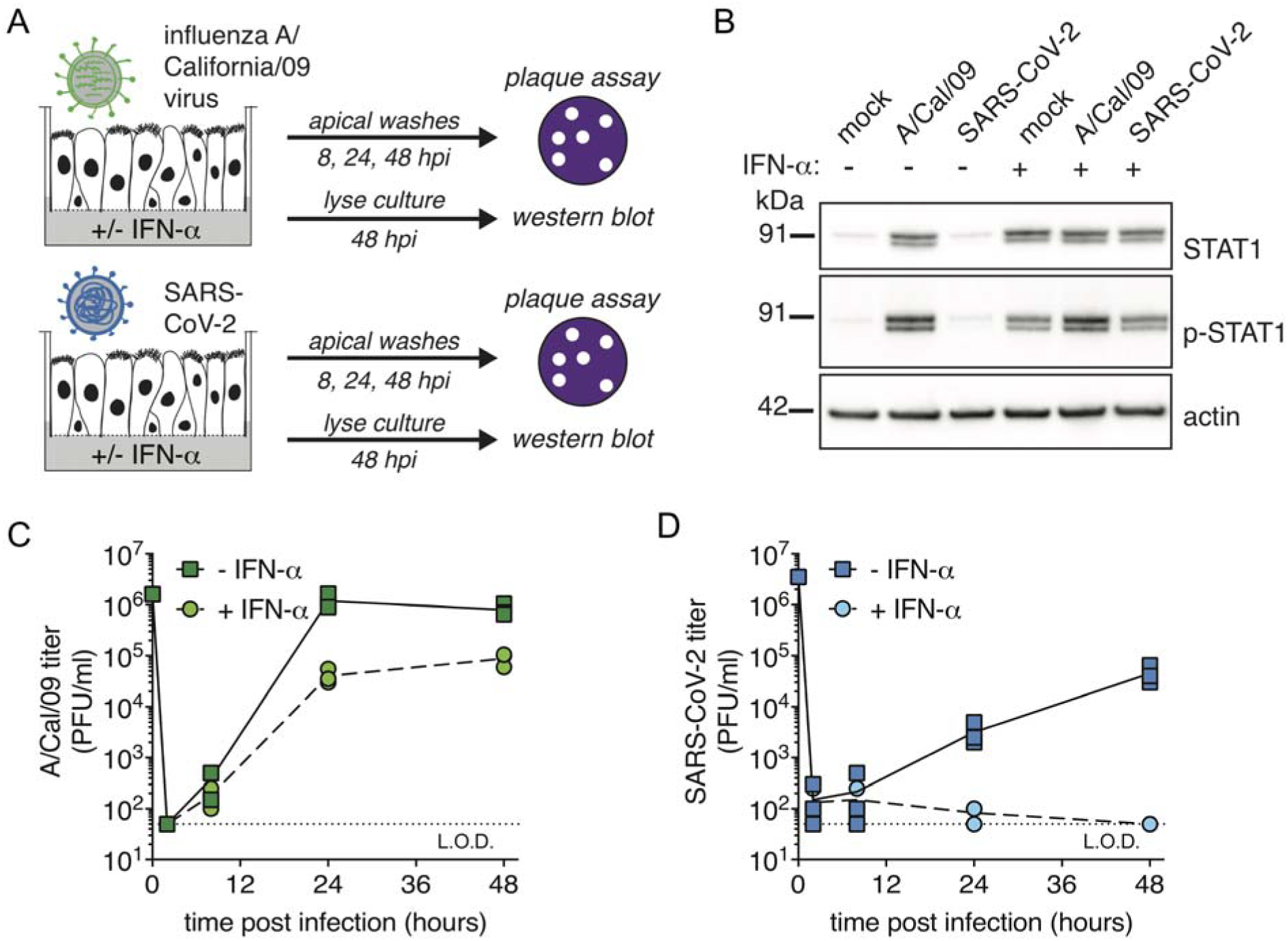
Differential IFN-I sensitivity and pSTAT1 phosphorylation following SARS-CoV-2 or influenza A virus on polarized human airway epithelial cultures (HAEC). A) HAEC were pretreated with 1000 U/mL IFN-α basolaterally for 2 hours prior to and during infection. Cultures were then infected apically with influenza A/California/09 H1N1 virus or SARS-CoV-2. Apical washes were collected at the indicated times and progeny titers determined by plaque assay on MDCK cells (influenza virus) or Vero E6 cells (SARS-CoV-2). At the 48 h endpoint, cultures were lysed for western blot analysis. B) Western for total STAT1 or phospho-STAT1 at 48 hpi. C) Influenza A virus titers by plaque assay on MDCK cells. 0 h, virus inoculate; 2 h, virus in third apical wash; 8, 24, 48 h, virus in apical washes at these time points. D) SARS-CoV-2 titers by plaque assay on Vero E6 cells. 0 h, virus inoculate; 2 h, virus in second apical wash; 8, 24, 48 h, virus in apical washes at these time points.

### SARS-CoV-2 impacted by IFN-I post treatment

Having established that SARS-CoV-2 cannot overcome a pre-induced IFN-I state, we next evaluated the impact of IFN-I treatment post infection. Vero cells were infected with SARS-CoV or SARS-CoV-2 at an MOI 0.01 and subsequently treated with 1000 U/mL IFN-α four hours post infection. For SARS-CoV, treatment post infection had no significant impact on viral yields either 24 or 48 hours post infection, consistent with findings from pretreatment experiments. In contrast, SARS-CoV-2 had a substantial 2-log_10_ reduction in viral titers at 24 hours post infection relative to control. However, by 48 hours, SARS-CoV-2 replication had achieved similar level to untreated controls indicating that post treatment had only a transient impact in Vero cells. We subsequently performed the post-treatment experiment utilizing Calu3 respiratory cells at an MOI 1. Similar to Vero cells, SARS-CoV infection post treatment resulted in no significant changes at 24 hours and has a modest decrease (∼0.5 log_10_) at 48 hours, illustrating its resistance to IFN-I. In contrast, IFN-I post-treatment of SARS-CoV-2 infection resulted in a >2-log_10_ reduction in viral titers at 24 hours and expanded at 48 hours post (∼4-log_10_ reduction). The results indicate that in Calu3 cells, SARS-CoV-2 is unable to prevent inhibitory effects of IFN-I even after establishing initial infection. Overall, the pre- and post-treatment data highlight distinct differences between SARS-CoV and SARS-CoV-2 in modulation of IFN-I pathways.

### Conservation of IFN-I antagonists across SARS-CoV and SARS-CoV-2

Previous work has established several key IFN-I antagonists in the SARS-CoV genome, including NSP1, NSP3, ORF3b, ORF6, and others (28). Considering SARS-CoV-2’s sensitivity to IFN-I, we next sought to evaluate conservation of IFN-I antagonist proteins encoded by SARS-CoV-2, SARS-CoV, and several bat SARS-like viruses including WIV16-CoV (29), SHC014-CoV (30), and HKU3.1-CoV (31). Using sequence analysis, we found several changes to SARS-CoV-2 that potentially contribute to IFN-I sensitivity (**Fig. 4**). For SARS-CoV structural proteins, including the nucleocapsid (N) and matrix (M) protein, a high degree of sequence homology (>90%AA identity) suggests that their reported IFN-I antagonism is likely maintained in SARS-CoV-2 and other SARS-like viruses. Similarly, the ORF1ab poly-protein retains high sequence identity in SARS-CoV-2 and several known IFN-I antagonists contained within the poly-protein (NSP1, NSP7, NSP14-16) are highly conserved relative to SARS-CoV. One notable exception is the large papain-like protease, NSP3, which is only 76% conserved between SARS-CoV and SARS-CoV-2. However, SARS-CoV-2 does maintain a deubiquitinating domain thought to confer IFN-I resistance (32). For SARS-CoV ORF3b, a 154 amino acid (AA) protein known to antagonize IFN-I responses by blocking IRF3 phosphorylation (33), sequence alignment indicates that the SARS-CoV-2 equivalent ORF3b contains a premature stop codon resulting in a truncated 24 AA protein. Similarly, HKU3.1-CoV also has a premature termination resulting in a predicted 39 AA protein. Both WIV16-CoV and SHC014-CoV, the most closely related bat viruses to SARS-CoV, encode longer 114 AA truncated protein with >99% homology with SARS-CoV ORF3b suggesting that IFN-I antagonism might be maintained in these specific group 2B CoV strains. In addition, SARS-CoV ORF6 has been shown to be an IFN-I antagonist that disrupts karyopherin-mediated transportation of transcription factors like STAT1 (33, 34). In contrast to ORF3b, all five surveyed group 2B CoVs maintain ORF6; however, SARS-CoV-2 had only 69% homology with SARS-CoV while the other three group 2B bat CoVs had >90% conservation. Importantly, SARS-CoV-2 has a two amino acid truncation in its ORF6; previous work has found that alanine substitution in this C-terminus of SARS-CoV ORF6 resulted in ablated antagonism (34). Together, the sequence homology analysis suggests that differences in NSP3, ORF3b, and/or ORF6 may be key drivers of SARS-CoV-2 IFN-I susceptibility.

**Figure 4.**
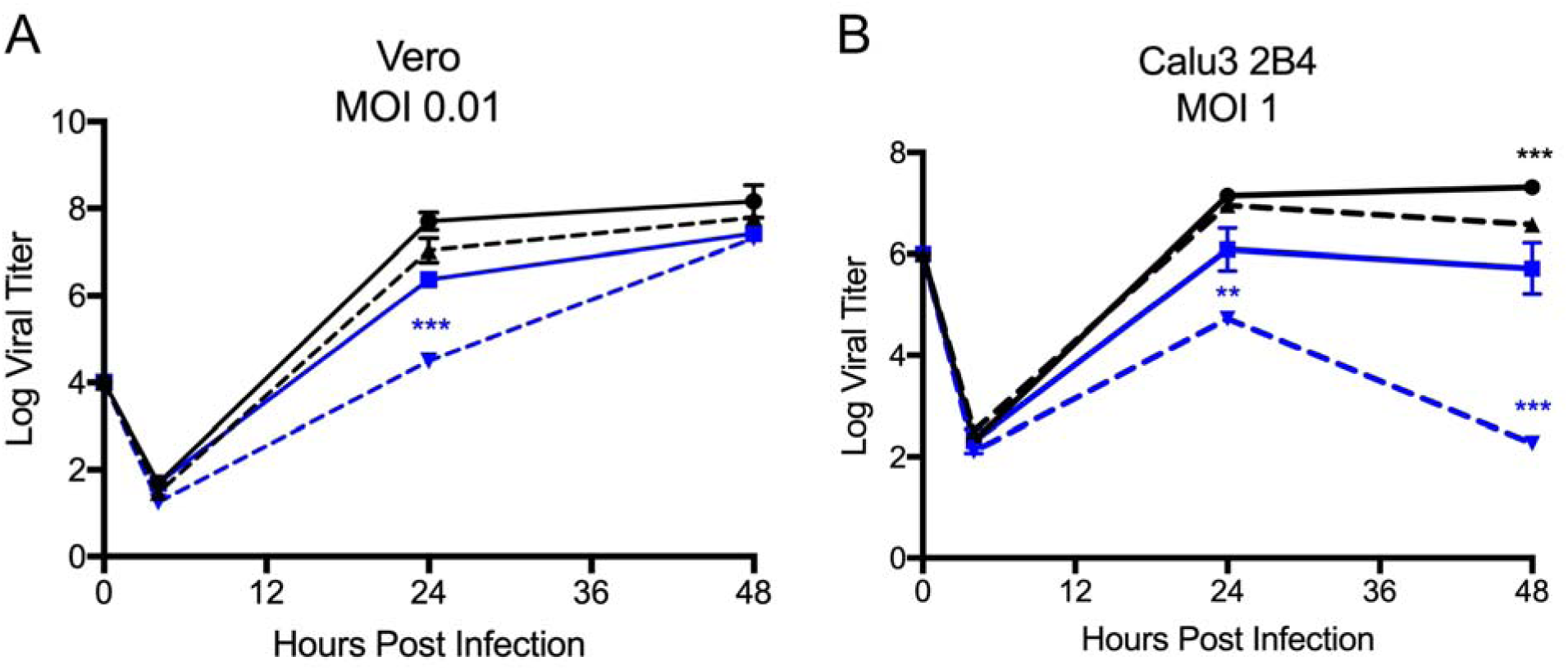
SARS-CoV-2 impacted by post IFN-I treatment. A-B) Vero E6 and Calu3 2B4 cells were infected with either SARS-CoV WT (black) or SARS-CoV-2 (blue) at an MOI 0.01 (A, Vero cells) or MOI 1 (B, Calu3 cells). Cells were subsequently treated with 1000 U/mL recombinant type I IFN (hashed line) or mock (solid line) for 4 hours following infection. Each point on the line graph represents the group mean, N ≥ 3. All error bars represent SD. The two tailed student’s t-test was used to determine P-values: **P <0.01 *** P < 0.001.

**Figure 5.**
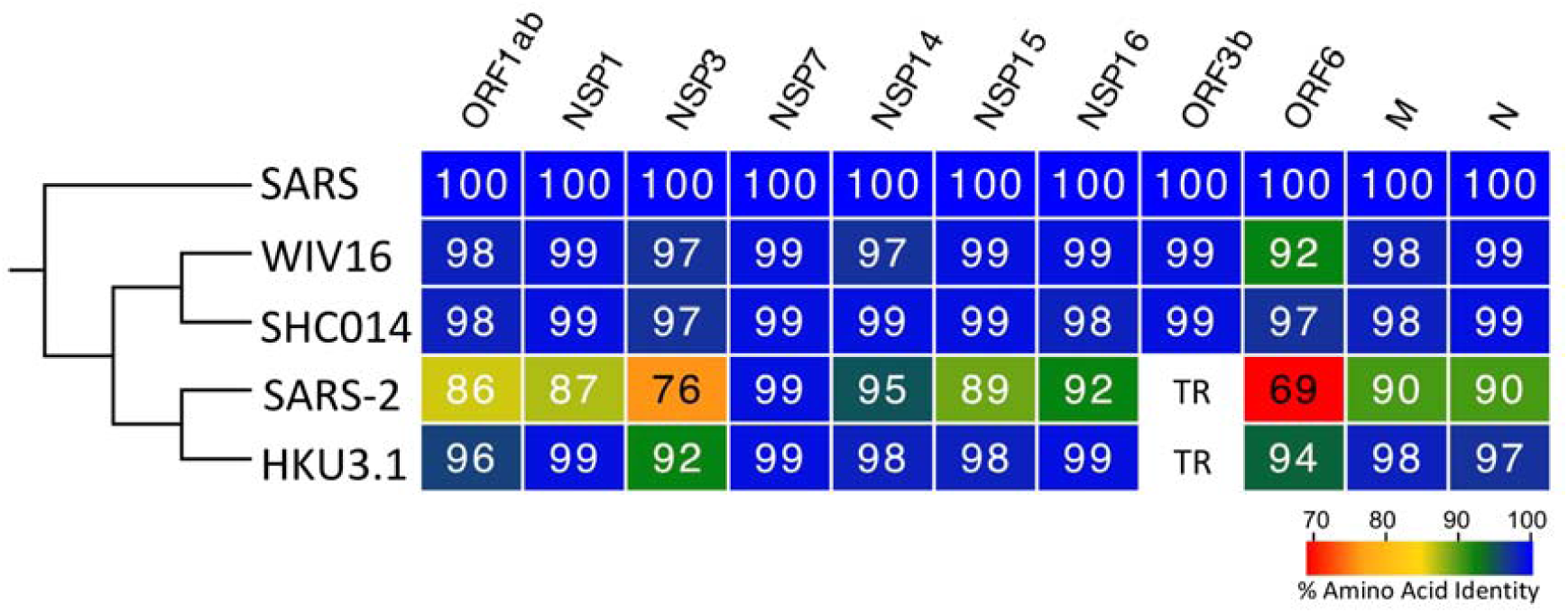
Conservation of SARS-CoV IFN antagonists. Viral protein sequences of the indicated viruses were aligned according to the bounds of the SARS-CoV open reading frames for each viral protein. Sequence identities were extracted from the alignments for each viral protein, and a heat map of percent sequence identity was constructed using EvolView (www.evolgenius.info/evolview) with SARS-CoV as the reference sequence. TR = truncated protein.

## Discussion

With the ongoing outbreak of COVID-19 caused by SARS-CoV-2, viral characterization remains a key factor in responding to the emergent novel virus. In this report, we describe differences in the IFN-I sensitivity between SARS-CoV-2 and the original SARS-CoV. While both viruses maintain similar replication in untreated Vero E6 cells, SARS-CoV-2 has a significant decrease in viral protein and replication following IFN-I pretreatment. The decreased SARS-CoV-2 replication correlates with phosphorylation of STAT1 and augmented ISG expression largely absent following SARS-CoV infection despite IFN-I pretreatment. Infection of IFN-I competent Calu3 2B4 cells resulted in reduced SARS-CoV-2 replication relative to SARS-CoV; IFN-I pretreatment also corresponded to increased sensitivity for SARS-CoV-2 in Calu3 cells. However, SARS-CoV-2 fails to induce IFN-I pathways during infection of unprimed polarized HAEC as compared to influenza A virus, suggesting a capacity to control IFN-I signaling. Yet, pretreatment nearly ablates replication of SARS-CoV-2 highlighting robust sensitivity to IFN-I pathways once activated. Finally, SARS-CoV-2, unlike SARS-CoV, responded to post-infection IFN-I treatment with reduced viral yields in both Vero and Calu3 cells. Analysis of viral proteins finds SARS-CoV-2 has several changes that potentially impact its capacity to modulate the IFN-I response, including loss of an equivalent ORF3b and a short truncation of ORF6, both known as IFN-I antagonists for SARS-CoV (33). Together, our results suggest SARS-CoV and SARS-CoV-2 have differences in their ability to antagonize the IFN-I response once initiated and that this may have major implications for COVID-19 disease and treatment.

With a similar genome organization and disease symptoms in humans, the SARS-CoV-2 outbreak has drawn insights from the closely related SARS-CoV. However, the differences in sensitivity to IFN-I pretreatment illustrate a clear distinction between the two CoVs. Coupled with a novel furin cleavage site (35), robust upper airway infection (8), and transmission prior to symptomatic disease (36), the differences between SARS-CoV and SARS-CoV-2 could prove important in disrupting the ongoing spread of COVID-19. For SARS-CoV, *in vitro* studies have consistently found that wild-type SARS-CoV is indifferent to IFN-I pretreatment (37, 38). Similarly, *in vivo* SARS-CoV studies have found that the loss of IFN-I signaling had no significant impact on disease (39), suggesting that this virus is not sensitive to the antiviral effects of IFN-I. However, more recent reports suggest that host genetic background may majorly influence this finding (40). For SARS-CoV-2, our results suggest that IFN-I pretreatment produces a 3 – 4 log_10_ drop in viral titer. This level of sensitivity is similar to MERS-CoV and suggests that the novel CoV lacks the same capacity to escape a primed IFN-I response as SARS-CoV (41, 42). Notably, the sensitivity to IFN-I does not completely ablate viral replication; unlike SARS-CoV 2’O methyl-transferase mutants (37), SARS-CoV-2 is able to replicate to low, detectable levels even in the presence of IFN-I. This finding could help explain positive test results in patients with minimal symptoms and the range of disease observed. In addition, while SARS-CoV-2 is sensitive to IFN-I pretreatment, both SARS-CoV and MERS-CoV employ effective means to disrupt virus recognition and downstream signaling until late during infection (25). While SARS-CoV-2 may employ a similar mechanism early during infection, STAT1 phosphorylation and reduced viral replication are observed in IFN-I primed and post-treatment conditions indicating that the novel CoV does not block IFN-I signaling as effectively as the original SARS-CoV.

For SARS-CoV-2, the sensitivity to IFN-I indicates a distinction from SARS-CoV and suggests differential host innate immune modulation between the viruses. The distinct ORF3b and truncation/changes in ORF6 could signal a reduced capacity of SARS-CoV-2 to interfere with IFN-I responses. For SARS-CoV ORF6, the N-terminal domain has been shown to have a clear role in its ability to disrupt karyopherin-mediated STAT1 transport (34); in turn, the loss or reduction of ORF6 function for SARS-CoV-2 would likely render it much more susceptible to IFN-I pretreatment as activated STAT1 has the capacity to enter the nucleus and induce ISGs and the antiviral response. In these studies, we have found that following IFN-I pretreatment, STAT1 phosphorylation is induced following SARS-CoV-2 infection. The increase in ISG proteins (STAT1, IFITM1) suggests that SARS-CoV-2 ORF6 does not effectively block nuclear transport as well as SARS-CoV ORF6. For SARS-CoV ORF3b, the viral protein has been shown to disrupt phosphorylation of IRF3, a key transcriptional factor in the production of IFN-I and the antiviral state (33). While its mechanism of action is not clear, the ORF3b absence in SARS-CoV-2 infection likely impacts its ability to inhibit the IFN-I response and eventual STAT1 activation. Similarly, while NSP3 deubiquitinating domain remains intact, SARS-CoV-2 has a 24 AA insertion upstream of this deubiquitinating domain that could potentially alter that function (32). While other antagonists are maintained with high levels of conservation (>90%), single point mutations in key locations could modify function and contribute to increased IFN-I sensitivity. Overall, the sequence analysis suggests that differences between SARS-CoV and SARS-CoV-2 viral proteins may drive attenuation in the context of IFN-I pretreatment.

The increased sensitivity of SARS-CoV-2 suggests utility in treatment using IFN-I. While IFN-I has been used in response to chronic viral infection (43), previous examination of SARS-CoV cases found inconclusive effect for IFN-I treatment (44). However, the findings from the SARS-CoV outbreak were complicated by combination therapy of IFN-I with other treatments including ribavirin/steroids and lack of a regimented protocol. While IFN-I has been utilized to treat MERS-CoV infected patients, no conclusive data yet exists to determine efficacy (45). Yet, *in vivo* studies with MERS-CoV have found that early induction with IFN-I can be protective in mice (46); importantly, the same study found that late IFN-I activation can be detrimental for MERS-CoV disease (46). Similarly, early reports have described treatments using IFN-I in combination for SARS-CoV-2 infection; yet the efficacy of these treatments and the parameters of their use are not known (47). Overall, sensitivity data suggest that IFN-I treatment may have utility for treating SARS-CoV-2 if the appropriate parameters can be determined. In addition, use of type III IFN, which is predicted to have utility in the respiratory tract, could offer another means for effective treatment for SARS-CoV-2.

In addition to treatment, the sensitivity to IFN-I may also have implications for animal model development. For SARS-CoV, mouse models that recapitulate human disease were developed through virus passage in immune competent mice (48). Similarly, mouse models for MERS-CoV required adaptation in mice that had genetic modifications of their dipeptidyl-peptidase 4 (DPP4), the receptor for MERS-CoV (49, 50). However, each of these MERS-CoV mouse models still retained full immune capacity. In contrast, SARS-CoV-2 sensitivity to IFN-I may signal the need to use an immune deficient model to develop relevant disease. While initial work has suggested incompatibility to SARS-CoV-2 infection in mice based on receptor usage (8), the IFN-I response may be a second major barrier that needs to be overcome. Similar to the emergent Zika virus outbreak, the use of IFN-I receptor knockout mice or IFN-I receptor blocking antibody may be necessary to develop a useful SARS-CoV-2 animal models for therapeutic testing (51).

Overall, our results indicate that SARS-CoV-2 has a much higher sensitivity to type I IFN than the previously emergent SARS-CoV. This augmented type I IFN sensitivity is likely due to changes in viral proteins between the two epidemic CoV strains. Moving forward, these data could provide important insights for both the treatment of SARS-CoV-2 as well as developing novel animal models of disease. In this ongoing outbreak, the results also highlight a distinction between the highly related viruses and suggest insights from SARS-CoV must be verified for SARS-CoV-2 infection and disease.

## Methods

### Viruses and cells

SARS-CoV-2 USA-WA1/2020 was provided by the World Reference Center for Emerging Viruses and Arboviruses (WRCEVA) or BEI Resources and was originally obtained from the USA Centers of Disease Control as described (52). SARS-CoV-2 and mouse-adapted recombinant SARS-CoV (MA15) (48) were titrated and propagated on Vero E6 cells, grown in DMEM with 5% fetal bovine serum and 1% antibiotic/antimytotic (Gibco). Calu3 2B4 cells were grown in DMEM with 10% defined fetal bovine serum, 1% sodium pyruvate (Gibco), and 1% antibiotic/antimitotic (Gibco). Standard plaque assays were used for SARS-CoV and SARS-CoV-2 (26, 53). All experiments involving infectious virus were conducted at the University of Texas Medical Branch (Galveston, TX) or New York University School of Medicine (New York City, NY) in approved biosafety level 3 (BSL) laboratories with routine medical monitoring of staff.

### Infection and type I IFN pre- and post-treatment

Viral replication studes in Vero E6 and Calu3 2B4 cells were performed as previously described (37, 54). Briefly, cells were washed two times with PBS and inoculated with SARS-CoV or SARS-CoV-2 at a multiplicity of infection (MOI) 0.01 for 60 minutes at 37 °C. Following inoculation, cells were washed 3 times, and fresh media was added to signify time 0. Three or more biological replicates were harvested at each described time. No blinding was used in any sample collections, nor were samples randomized. For type I IFN pretreatment, experiments were completed as previously described (37). Briefly, Vero E6 cells were incubated with 1000 U/mL of recombinant type I IFN-α (PBL Assay Sciences) 18 hours prior to infection (37). Cells were infected as described above and type I IFN was not added back after infection.

### Generation of polarized human airway epithelial cultures (HAEC)

hTert-immortalized human normal airway tracheobronchial epithelial cells, BCi.NS1.1 (55) were maintained in ExPlus growth media (StemCell). To generate HAEC, BCi.NS1.1 were plated (7.5E4 cells/well) on rat-tail collagen type 1-coated permeable transwell membrane supports (6.5mm; Corning Inc), immersed in ExPlus growth media in both the apical and basal chamber. Upon reaching confluency, media in the apical chamber was removed (airlift), and media in the basal chamber was changed to Pneumacult ALI maintenance media (StemCell). Pneumacult ALI maintenance media was changed every two days for approximately 6 weeks to form differentiated, polarized cultures that resemble in vivo pseudostratified mucociliary epithelium.

### IFN-I treatment and infection of HAEC

For interferon pretreatment of human airway epithelial cultures, 1000 U/mL of universal recombinant interferon-alpha was added to the basolateral chamber 2 h prior to viral infections. For viral infections, cultures were washed apically with 50 *µ*l of prewarmed PBS three times for 30 min at 37°C. Virus inoculates (1.3E6 PFU for influenza A/California/07/2009 (H1N1) virus and 1.35E5 PFU for SARS-CoV-2 USA/WA/1/2020 in 50 *µ*l PBS Mg/Ca) were added apically for two hours. Inoculates were then removed and cultures were washed three times with PBS, the third wash was collected and then the cultures were incubated for the indicated times. Progeny virus was collected in apical washes with 50 *µ*l PBS Mg/Ca for 30 min at 8, 24 and 48 hpi (endpoint). At endpoint, the membranes were collected and lysed in 300 *µ*l LDS lysis buffer (Life Technologies). Protein levels were measured by western blot. Briefly, 5 *µ*l of cell lysates were loaded into a SDS-PAGE gel and transferred to a nitrocellulose membrane. The membranes were blocked for 1 h at room temperature with 5% skim milk and then incubated overnight at 4°C with appropriate primary antibody dilution: anti-STAT1 (1:1000, Cell Signaling), anti-pSTAT1 (1:1000, Cell Signaling) or anti-beta actin (1:5000, Invitrogen). Then, the membranes were incubated with the recommended dilution of HRP-conjugated secondary antibody (goat anti-rabbit IgG HRP 1:10000, Invitrogen; goat anti-mouse IgG HRP 1:10000, Invitrogen) at room temperature for 1 h. Super Signal West Dura Kit (Thermo Scientific) was used for signal development according to manufacturer’s recommendations and the chemiluminescence was detected with a ChemiDoc Imager (BioRad).

### Phylogenetic Tree and Sequence Identity Heat Map

Heat maps were constructed from a set of representative group 2B coronaviruses by using alignment data paired with neighbor-joining phylogenetic trees built in Geneious (v.9.1.5). Sequence identity was visualized using EvolView (http://evolgenius.info/) and utilized SARS-CoV Urbani as the reference sequence. Tree shows the degree of genetic similarity of SARS-CoV-2 and SARS-CoV across selected group 2B coronaviruses.

### Immunoblot Analysis and Antibodies

Viral and host protein analysis were evaluated as previously described (52, 56). Briefly, cell lysates were resolved on 7.5% Mini-PROTEAN TGX SDS-PAGE gels and then transferred to polyvinylidene difluoride (PVDF) membranes using a Trans-Blot Turbo transfer system (Bio-Rad). Membranes were blocked with 5% (w/v) non-fat dry milk in TBST (TBS with 0.1% (v/v) Tween-20) for 1 hr, and then probed with the indicated primary antibody in 3% (w/v) BSA in TBST at 4°C overnight. Following overnight incubation, membranes were probed with the following secondary antibodies in 5% (w/v) non-fat dry milk in TBST for 1 hr at room temperature: anti-rabbit or anti-mouse IgG-HRP conjugated antibody from sheep (both 1:10,000 GE Healthcare). Proteins were visualized using ECL or SuperSignal West Femto chemiluminescence reagents (Pierce) and detected by autoradiography. The following primary antibodies were used: anti-pSTAT1 (Y701) (1:1000 9171L Cell Signaling Technologies), anti-STAT1 D1K9Y (1:1000 14994P Cell Signaling Technologies), anti-IFITM1 (1:1000 PA5-20989 Invitrogen), anti-SARS-CoV/CoV-2 Spike 1A9 (1:1000 GTX632604 GeneTex), and anti-β-Actin (1:1000 ab8227 Abcam).

### Statistical analysis

All statistical comparisons in this manuscript involved the comparison between 2 groups, SARS-CoV or SARS-CoV-2 infected groups under equivalent conditions. Thus, significant differences in viral titer were determined by the unpaired two-tailed student’s T-Test.

## Acknowledgements

Research was supported by grants from NIA and NIAID of the NIH (U19AI100625 and R00AG049092 to VDM; R24AI120942 to WRCEVA; R01AI134907 to RR; 1R01AI143639-01 and 1R21AI139374-01 to MD; Jan Vilcek/David Goldfarb Fellowship Endowment Funds to AMVJ; and T32 AI007526 to AH). Research was also supported by STARs Award provided by the University of Texas System to VDM and trainee funding provided by the McLaughlin Fellowship Fund at UTMB.

## Notes

### Competing Interest Statement

The authors have declared no competing interest.

### Summary of Updates

This manuscript adds additional data for pretreatment of Calu3 with IFN-I. Also compares immune stimulation/pretreatment in HAEC relative to influenza A virus Evaluates efficacy of IFN-I treatment post infection.

